# Quantitative study of alpha-synuclein prion-like spreading in fully oriented reconstructed neural networks reveals non-synaptic dissemination of seeding aggregates

**DOI:** 10.1101/2021.10.06.463379

**Authors:** Josquin Courte, Ngoc Anh Le, Luc Bousset, Ronald Melki, Catherine Villard, Jean-Michel Peyrin

**Author notes:** **Corresponding author** JM Peyrin.

## Abstract

The trans-neuronal spread of protein aggregates in a prion-like manner underlies the progression of neuronal lesions in the brain of patients with synucleinopathies such as Parkinson’s disease. Despite being studied actively, the mechanisms of alpha-synuclein (aSyn) aggregates propagation remain poorly understood. This hinders the development of therapeutic approaches aiming at preventing the spatial progression of intracellular inclusions in neural networks. To assess the role of synaptic structures and neuron characteristics in the transfer efficiency of aggregates with seeding propensity, we developed a novel microfluidic culture system which allows for the first time to reconstruct in vitro fully oriented and synaptically connected neural networks. This is achieved by filtering axonal growth with unidirectional “axon valves” microchannels. We exposed the presynaptic compartment of reconstructed networks to well characterized human aSyn aggregates differing in size: Fibrils and Oligomers. Both aggregates were transferred to postsynaptic neurons through active axonal transport, albeit with poor efficiency. By manipulating network maturity, we compared the transfer rate of aggregates in networks with distinct levels of synaptic connectivity. Surprisingly, we found that transfer efficiency was lower in mature networks with higher synaptic connectivity. We then investigated the seeding efficiency of endogenous aSyn in the postsynaptic population. We found that exposure to Fibrils, and not Oligomers, resulted in low efficiency trans-neuronal seeding which was restricted to postsynaptic axons. Finally, we assessed the impact of neuron characteristics and aSyn expression on the propagation of aSyn aggregates. By reconstructing chimeric networks, we found that neuron characteristics, such as the brain region from which they originate or aSyn expression levels, did not significantly impact aggregates transfer, and observed no trans-neuronal seeding where the presynaptic population did not express aSyn. Overall, we demonstrate that this novel platform uniquely allows the quantitative interrogation of original aspects of the trans-neuronal propagation of seeding pathogenic entities.

## Background

Chronic neurodegenerative diseases (ND) such as synucleinopathies are characterized by slow and progressive neuronal dysfunctions that occur over years in patients. It is now well established that a subset of neurodegenerative syndromes involves the progressive accumulation of abnormal protein aggregates in neurons [41]. Alpha-synuclein (aSyn) aggregation, which is associated Lewy Bodies (LB), is one of the major hallmarks of synucleinopathies and is hypothesized to play an important role in the etiology of Parkinson’s disease (PD) related neuronal dysfunction [13]. Recent evidences from neuropathological studies shows that LB accumulation progresses both spatially and temporally in the brains of affected patients and follows a rostro-caudal stereotyped pattern [48]. Somewhat akin to the pathogenic form of the prion protein, aggregated aSyn spreads from cell to cell and amplifies by seeding the aggregation of its soluble counterpart [17, 27, 28, 33, 39]. Importantly, protein aggregates build up as structural polymorphic structures holding specific biological activities reminiscent of prion strains [22, 32, 45, 49, 55]. To date, research in PD mainly focused on linking clinical phenotypes to cell intrinsic genetic and cellular alterations. This led to the demonstration that specific protein conformers account for progressive neuronal dysfunction and/or death in both sporadic or genetic age-related ND. Yet both the molecular and cellular mechanisms involved in the spatial spreading of aSyn aggregates between neurons and along neural pathways remains poorly understood. Several studies suggests that aSyn aggregates can be conveyed in extracellular medium and spread to neighboring cells naked, trough small extracellular vesicles or tunneling nanotubes in immortalized cell lines and primary neuronal cultures [1, 6, 11, 16, 56]. The respective contribution of these mechanisms to aSyn aggregates remain unclear. Because sequentially affected regions in the brain are synaptically connected, synapses are postulated to play a role in aSyn aggregates trans-neuronal spreading [10, 43]. Indeed, synapses, which tightly link neuronal membranes, harbor putative receptors of aSyn aggregates [47]. Moreover, synapse are hotspots of membrane remodeling [15] and contain most of the pool of soluble -and vulnerable to seeding-aSyn in their presynaptic moiety. Yet, their role in aSyn transfer has not been clearly evaluated: synaptic connectivity alone does not predict the pattern of aggregation spreading in the brain [20, 21, 50]. Whether or not synapses are permissive for protein diffusion or filters aggregates according to their size and how much aSyn aggregates can diffuse from one neuron to the other, are pending questions.

While these questions are barely testable in vivo, microfluidic approaches, which allow in vitro reconstruction of neural networks, may prove helpful to address these issues [8, 11, 14]. Here, we describe a new microfluidic design which allows full control of axonal outgrowth direction and therefore reconstruction of fully oriented binary neuronal networks. We selectively challenged presynaptic neurons with fluorescently labeled aSyn pre-formed fibrils (PFF) and lower molecular weight oligomers and quantitively studied the 2 distinct steps that underlies their prion-like spreading in the brain: 1) trans-neuronal dissemination of aSyn aggregates, and 2) seed-induced aSyn nucleation in post-synaptic neurons. While aSyn oligomers efficiently spread in networks, we observed that seeding competent aSyn PFF have relatively low trans-neuronal dissemination capacities with 1% of presynaptic aggregates spread to second order neurons in 3 days. This translates in delayed post synaptic aSyn nucleation, which occurred in the distal, axonal parts of the post synaptic neurons. Moreover, by reconstructing mosaic networks we demonstrate that the process is not modified by the characteristics of the emitting neuron nor its aSyn expression level. These results provide key kinetics parameters of the 2 critical prion-like steps which kinetics are likely to shape the overall dissemination in neural networks.

## Methods

### Microfluidic culture chips

A detailed version of the protocol used for the fabrication of SU-8 master mold, resin mold and culture chips can be found in [8]. Briefly, microchannels and culture compartments were drawn using QCAD. The dimensions of the chips with axon valves microchannels are indicated in Supp Fig 1. In experiments involving chips with straight microchannels, all dimensions were identical, except for the 10 axon valves microchannels that were replaced by 10 straight 10 μm wide microchannels. Photo-masks engraved with the designs were ordered from Selba. Photosensitive negative SU-8 resin (MicroChem) was spin-coated to the desired height on a silicon wafer (Prolog Semicor) and exposed to UV light through a photo-mask encoding the culture chip outlines using an MJB4 mask aligner (Suss). Unexposed SU-8 was washed away. These steps were repeated twice, once with a low viscosity resin for 3 μm high microchannels, and once with a higher viscosity resin for 50 μm high culture compartments. The height of the mold was then checked with a stylus profiler (Dektak), and microstructures integrity was verified with an optical microscope (Leica). The resulting mold was used to cast polydimethylsiloxane (PDMS Sylgard, Ellsworth Adhesives) mixed at a 10:1 w:w ratio of base to curing agent. PDMS was cured 3 h minimum at 70 °C. For high throughput chip production, a sturdier epoxy resin mold was used. The resulting chips were unmolded, and inlets were punched with a surgical biopsy punch (4 mm diameter) at both extremities of each compartment. The PDMS blocks were bonded to 130-160 μm thick glass coverslips (Fisher Scientific 11767065) after plasma surface treatment with an Atto plasma cleaner (Diener). Culture compartments were then immediately filled with deionized water. Devices were placed in individual Petri dishes for easier handling, and each of the resulting culture system was sterilized by exposure to a UV lamp for 30 min. The day before cells seeding, culture devices were coated with Phosphate Buffered Saline (PBS, Thermo Fisher 14190169) containing 10 μg/mL poly-D-lysine (Sigma P7280), introduced in one well per culture compartment. Devices were then incubated overnight in a 37 °C humidified atmosphere. 4-6 h before cells seeding, wells were emptied, then filled in the same fashion with PBS containing 2.5-5 μg/mL laminin (Sigma L2020).

### Primary neuronal cultures

Animal care was conducted in accordance with standard ethical guidelines (U.S. National Institutes of Health publication no. 85–24, revised 1985, and European Committee Guidelines on the Care and Use of Laboratory Animals) and the local, IBPS and UPMC, ethics committee approved the experiments (in agreement with the standard ethical guidelines of the CNRS “Formation à l′Expérimentation Animale” and were approved by the “C2EA -05 Comité d’éthique en experimentation animale Charles Darwin”).

Hippocampal (Hip) and cortical (Cx) regions were micro-dissected from embryos from pregnant mice at embryonic day 16 (E16). Wild type (WT, Hip^SNCA +/+^) cultures were prepared from inbred Swiss mice (Janvier). Hip cultures knockout for aSyn (Hip^SNCA -/-^) were prepared from inbred C57bl/6Jolahsd mice (Envigo). Hip cultures expressing membrane associated tandem Tomato (mtdTomato, Hip^mTmG +/-^) or not (Hip^mTmG -/-^) were obtained by cross-breeding C57bl/6J *mTmG +/-* (adapted from [31]) males with C57bl/6J *mTmG -/-* females. A fluorescent protein flashlight (Nightsea DFP-1) was used to distinguish *mTmG +/-* from *mTmG -/-* embryos.

Cx and Hip microstructures were washed in Gey’s Balanced Saline Solution (Sigma G9779) and chemically digested with papain (Sigma 76220) diluted (7.5 units/mL for Cx, 15 units/mL for Hip) in Dulbecco’s Modified Eagle Medium (DMEM, Thermofisher 31966 021) at 37 °C. After digestion was interrupted by adding 10 % Fetal Bovine Serum (FBS, GE Healthcare), supernatant was replaced with DMEM containing 40 units/mL of DNase type IV (Sigma D5025). Physical cell dissociation was performed by gently flowing microstructures 10 times through a 1 mL micropipette tip. Cell suspension was centrifuged (80 g, 8 min) and resuspended with DMEM. Cells concentration was assessed in a Malassez counting device. Cells were centrifuged again (80 g, 8 min) and suspended at the desired concentrations in culture medium (Neurobasal medium (Thermofisher 21103049) supplemented with 1 % GlutaMAX (Thermofisher 35050), 1 % B27 Supplement (Thermofisher 17504-044)) supplemented with 10 % FBS. The central compartment of microfluidic devices was filled with culture medium before cells seeding, to prevent cells from flowing through microchannels during cells seeding. 1.5 μL of cells suspension at 40 and 10 million cells/ml were introduced in the presynaptic and postsynaptic compartments, respectively, and were left to adhere for 15 min. Culture wells were filled with culture medium supplemented with 10 % FBS. Water containing 0.5 % ethylenediaminetetraacetic acid (EDTA, Sigma EDS) was added in the Petri dish, around the chip, to limit evaporation. Culture devices were then placed in a humidified atmosphere at 37 °C and 5 % carbon dioxyde. Culture medium supplemented with 10 % FBS was completely replaced 24 h after cell seeding by FBS-free culture medium. Half of the medium was replaced every week with fresh FBS-free culture medium.

### Immunostaining

For all steps, one inlet by compartment was filled to ensure solution flow though the culture compartment. Cultures were fixed at room temperature for 10 min in PBS containing 4 % paraformaldehyde (Electron Microscopy Science 15714S) and 4 % sucrose (Sigma S9378). Membrane permeabilization and saturation were performed simultaneously at room temperature for 30 min in PBS containing 0.2 % Triton X-100 (Sigma X100) and 1 % bovine serum albumin (BSA, Thermofisher 15561020). Primary and secondary antibodies were diluted in PBS 1 % BSA and were respectively incubated 16 h at 4 °C and 1 h at room temperature. A list of antibodies used in this study can be found in Supplementary Table 1.

### Image acquisition

Epifluorescence imaging was performed using a DMi8 microscope (Leica) fitted with a pE-4000 LED light source (CoolLED) and a sCMOS Flash 4.0 camera (Hamamatsu). A 40X NA 0.8 objective was used for high magnification imaging, and a 10X NA 0.4 objective for low magnification imaging.

Confocal imaging was performed with a SP5 set-up (Leica) fitted with a 63X NA 1.4 objective.

### aSyn aggregates generation

Mouse (m) or human (h) aSyn proteins were produces as described in (Ghee et al 2005 FEBS J). Human aSyn Oligomers (hOlig) were obtained as described in [35]. Mouse and human fibrils (mFib and hFib, respectively) were assembled as described in [4]. The nature of aSyn assemblies was assessed by Transmission Electron Microscopy (TEM) after adsorption of the aggregates onto carbon-coated 200 mesh grids and negative staining with 1% uranyl acetate using a Jeol 1400 transmission electron microscope. The images were recorded with a Gatan Orius CCD camera (Gatan, Pleasanton, CA, USA). aSyn oligomers and fibrils were fluorescently labeled in PBS buffer with ATTO 647 sulfo-NHS dye at a dye-protein ratio of 1:1 following manufacturer recommendation (ATTO-Tech, Gmbh) and as previously described [47]. Fibrillar aSyn was fragmented by sonication for 20 min in 2-ml Eppendorf tubes in a Vial Tweeter powered by an ultrasonic processor UIS250v (250 W, 2.4 kHz; Hielscher Ultrasonic, Teltow, Germany) to generate fibrillar particles with an average size 42-52 nm as assessed by TEM analysis. Aliquots (5μl at 350 μM, concentration of monomers) of aSyn aggregates were flash frozen in liquid nitrogen and stored at -80 °C. Samples were thawed in a 37 °C water bath on the day of culture treatment.

### Spiking of neuronal cultures with aSyn aggregates

Cultures in microfluidic devices were first selected on the basis of cellular integrity in both compartments, and robust axonal outgrowth from presynaptic to postsynaptic compartment. Treatment of the presynaptic compartment with hFib, hOlig of mFib was performed at DIV13-14 or 20-21. Culture wells were completely emptied, and inlets of the postsynaptic, central and presynaptic compartments were filled sequentially, respectively with 45, 35 and 10 μL of medium. Postsynaptic and central inlets were filled with fresh culture medium, and presynaptic inlets were filled with culture medium containing or not 500 nM of freshly thawed aggregates, thoroughly mixed with culture medium by pipetting up and down 30 times. To ensure that presynaptic neurons were homogeneously exposed to aggregates, 2.5 μL of the bottom inlet were transferred to the top inlet. 24 h after treatment, the inlets of the presynaptic compartment were emptied, and neurons were rinsed one time by filling one well with 15 μL fresh medium, waiting 1 min and then emptying both inlets. Both presynaptic inlets were then filled with 15 μL of fresh medium.

### Image analysis and quality control before analysis

Image analysis was performed using custom image analysis pipelines for Fiji version 2.0.0 [44]. The main steps of each analysis are explained below.

A first time before treatment, and a second time before analysis, cultures were selected on the basis of cell survival and robust axonal outgrowth to the second compartment. These parameters were determined from the Phase channel. In our experimental timeframe, we did not observe a lower survival in cultures treated with 500 nM hOlig, hFib or mFib in comparison to control ones.

Aggregates associated fluorescence was shown in all figures using the look-up table (LUT) Turbo, obtained from https://github.com/cleterrier/ChrisLUTs.

### Analysis of axonal filtration

Crossing mtdTomato positive axons in Hip^mTmG +/-^>Hip^mTmG -/-^ and Hip^mTmG -/-^>Hip^mTmG +/-^ networks were manually counted at the exit of microchannels in presynaptic or postsynaptic compartments from epifluorescence microscopy 10X fields.

### Analysis of synaptic density

Cultures were stained with Bassoon, MAP2 and TUJ1. aSyn staining was additionally performed in Hip^SNCA+/+^>Hip^SNCA-/-^ networks. Confocal Z-stacks were acquired with a 0.42 μm z step and 0.24×0.24 μm pixels. Z-stacks were then submitted to maximum intensity projection. The Bassoon channel was pretreated with a 5 pixel radius Subtract Background, while the aSyn channel was pretreated with a 20 pixel radius Subtract Background. Fields to be analyzed were selected on the basis of low dendrites density and high axonal density. Isolated dendrites traces were drawn from the MAP2 channel using the Simple Neurite Tracer plugin [25] on Fiji. Traces were used to determine neurite length, and were dilated to 3μm width regions of interest (ROIs). ROIs were then used to segment the sections of the Bassoon channel to be analyzed. The Trainable WEKA Segmentation plugin [3] on Fiji was trained to recognize Bassoon foci, and applied to all Bassoon channels. Probability maps were binarized. Particles larger than 1 pixel were numbered with Analyze Particles to generate individual Bassoon ROIs. For determining the density of aSyn+ synapses, the mean aSyn signal was measured in each Bassoon ROI. Synapses were then classified as aSyn+ or aSyn-by an arbitrary threshold on the aSyn mean intensity. A minimum of 650 μm of dendrites from at least 4 separate fields were considered in each individual culture compartment.

### Analysis of aSyn aggregates fluorescence in somas

For analyses involving exogenous fluorescent aggregates, 5 frames were taken in Z with a step of 2 μm, and analysis was performed on the maximum intensity Z-project of these frames. Cultures had to be imaged live, as aggregates associated fluorescence decreased drastically after fixation. Thus, neuronal somas had to be detected from the Phase channel. ROIs circling neuronal somas were automatically detected from the Phase channel using a modified version of the region-convoluted neural network (R-CNN) described in [18], available at the following URL: https://github.com/matterport/Mask_RCNN. The algorithm was trained over 150 iterations using 4000 ROIs of neuronal somas manually traced in 350 fields in both presynaptic and postsynaptic compartments. Parameters were adjusted to the size of the desired ROIs. Automatically generated ROIs were then manually validated and corrected before segmentation of fluorescence channels. The mean aggregates associated fluorescence was measured in each ROI. Results were normalized by the mean fluorescence in untreated cultures imaged during the same microscopy session. A minimum of 40 somas from at least 4 separate fields were considered in each individual culture compartment.

### Analysis of aSyn aggregates fluorescence in axons

Individual axon tracts in the central channel of culture devices were manually traced using the Phase channel. Analysis of the fluorescence channel was then performed in a 1 μm wide ROI surrounding manually traced axonal tracts. Results were normalized by the mean fluorescence in untreated cultures imaged during the same microscopy session. A minimum of 10 axons were considered in each individual culture compartment.

### Calculation of Transmitted Excess Signal

Transmitted Excess Signal (TES) in individual experiments was computed using the following equations:

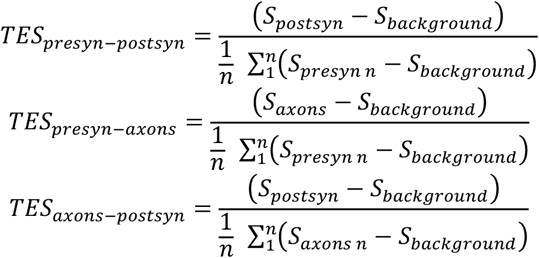

With *S*_*i*_ as the average of normalized signals in compartment *i* in an individual experiment, *S*_*background*_ as 1 (because ctrl values cluster around 1 after normalization), and *n* the total number of experiments for which signal in the presyn compartment was obtained.

### Distribution of aSyn aggregates associated fluorescence in individual somas

Distributions of signal in 500 randomly picked individual somas per condition were obtained by plotting data with the distplot() function of the Seaborn package on Python 3 with 100 bins.

### Analysis of endogenous aSyn aggregates

S129 phosphorylated aSyn (pSyn) and MAP2 were stained. Integrated signal densities were measured in the whole presynaptic and postsynaptic compartments, imaged by epifluorescence microscopy with a 10X objective. pSyn signal was normalized by MAP2 signal. Results were normalized by the pSyn/MAP2 ratio in untreated cultures imaged during the same microscopy session.

### Infection with adeno-associated viral vectors

Primary neuronal cortical-hippocampal network were cultured as previously described. At DIV5, hippocampal neurons of the postsyn chamber were transduced with AAV10 vectors (pssAAV-CBA-GFP-WPRE) with the dose of 10^8^ viral genomes/8.000 cells. At DIV8, the cell culture medium was replaced with fresh medium in order to eliminate the residual viral particles. Selective treatment of the postsyn compartment with viral vectors was performed in the same fashion than the spiking of the presyn compartment with aggregates described earlier. At DIV8, the presyn compartment was spiked with 500 nM mFib, as previously described. Cultures were then maintained for 22 days (17 days after AAV transfection at 14 days after aSyn exposure) at 37 ^0^C in 5 % CO_2_. They were then fixed and stained before imaging by confocal microscopy.

### Statistical analysis

Data analysis was performed with the Pandas package in Python 3. Statistical analysis and graphs generation were performed with GraphPad Prism version 8.4.2.

If not indicated otherwise, individual data points on graphs represent experimental means. Experimental means from at least 2 individual culture devices were used as individual data points for the purpose of statistical analysis.

First, the deviation from the normal (gaussian) distribution of each replicates in a condition was assessed with a Shapiro-Wilk’s test. Then, two-columns comparisons were performed with a t-test with Welch’s correction if both columns followed a normal distribution and with a Mann-Whitney test otherwise. Comparisons of more than two columns with one parameter varying were performed with a Brown-Forsythe ANOVA test followed by Dunnett’s T3 multiple comparisons test if all columns followed a normal distribution, and with Kruskal-Wallis test followed by Dunn’s multiple comparisons test otherwise. Comparisons of more than two columns with two parameters varying were performed with two-way ANOVA with the Geisser-Greenhouse correction followed by Tukey’s multiple comparisons test.

## Results

### Reconstruction of fully oriented neural networks in vitro

There is an unmet need for in vitro culture models dedicated to the study of pathogens propagation in neural networks. Indeed, most previous studies have taken advantage of neuronal microfluidic culture devices allowing axonal compartmentalization with straight axonal guiding microchannels, and these suffer from several pitfalls, detailed in the Discussion section.

We present here a compartmentalized culture system which satisfies the requirements for studying trans-neuronal propagation of protein particles with prion-like properties: unidirectional axonal growth and synaptic connectivity. Three compartments, 50μm in height, are linked by arrays of 3μm height axon-permissive microchannels (Fig 1a). While the two peripheral compartments, 1 mm wide, are dedicated to neuronal cultures, the central 50 μm wide compartment serves to observe axonal outgrowth and to improve fluidic isolation. We built on our previous approaches for guiding the direction of axonal growth [8, 34, 37] to design new asymmetric microchannels to build a device that does not allow axons from the postsynaptic chamber to grow to the presynaptic chamber, and took inspiration from the fluidic “Tesla” valves (US patent #US001329559) to name them “axon valves” (Fig 1b, Supp Fig 1). In the forward direction, axons are funneled by large openings (∼80 μm) into the microchannels maze which they can cross to the other side by following a “zig-zag” path (Fig 1b, green neuron). In the backward direction, the size of microchannels openings is much smaller (6 μm), and axons which penetrate the maze are submitted to a series of divergent Y junctions (Fig 1b, red neurons). As axons exhibit flexural rigidity, the path they are the most likely to follow sends them back to the compartment they came from, due to the presence of “arches-shaped” microchannels [37]. In addition, “dead-end” microchannels trap part of the axons, with the goal of preventing them from growing alongside the walls until they find one of the narrow openings. We will call the axon emitting compartment “presynaptic” (presyn), central compartment “axonal” and receiving compartment “postsynaptic” (postsyn) from now on. We further biased axonal growth by seeding the postsyn compartment with 4 times less neurons than the presyn compartment. It is noteworthy that, probably due to differences in trophic support, neuronal densities in presyn and postsyn compartments diverged by approximately 4 to 12 fold in later culture times (Supp Fig 2b).

**Figure 1.**
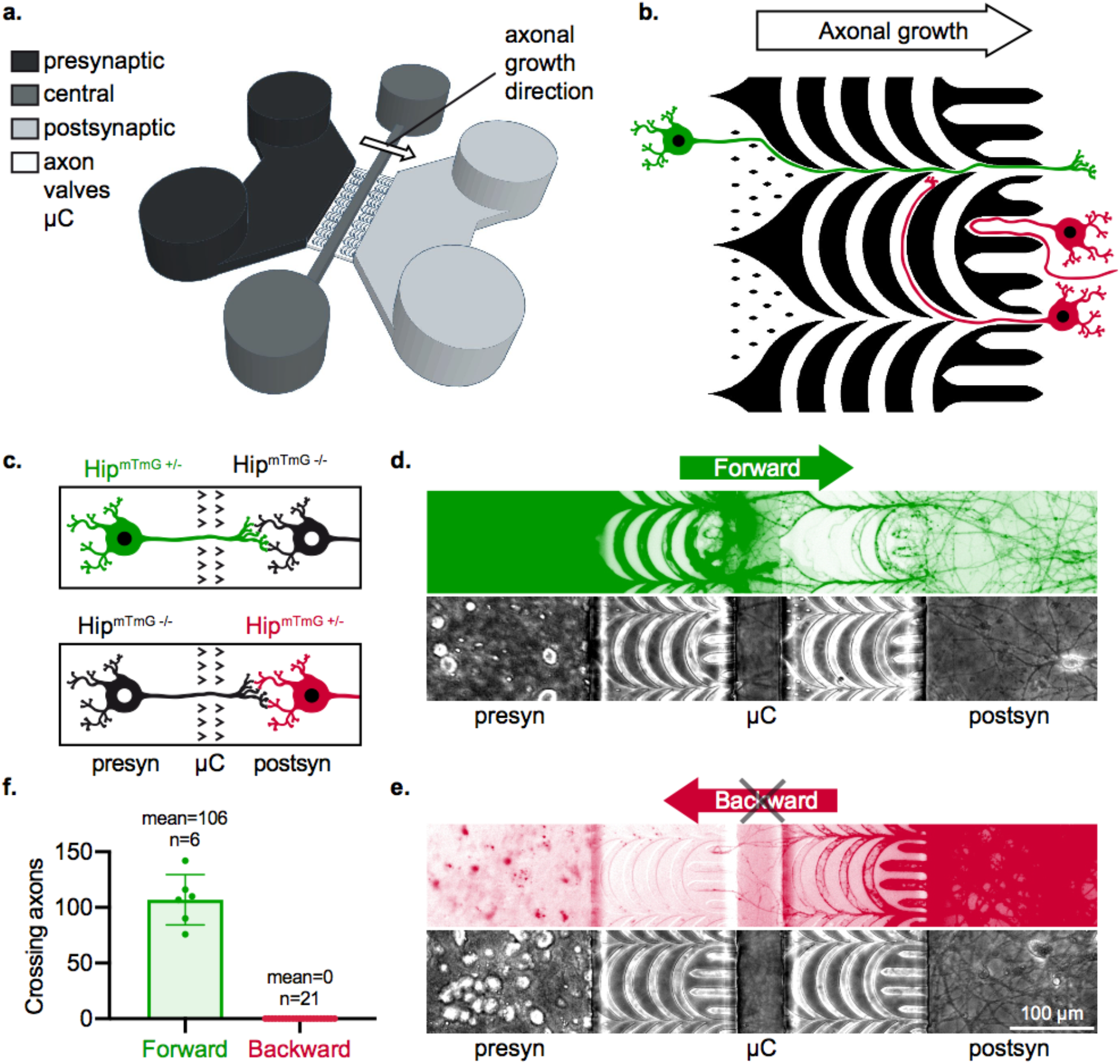
Reconstruction of fully oriented neural networks of chosen composition *in vitro*. **(a)** Schematic representation of the hollow parts of the microfluidic culture device for oriented neural networks reconstruction. Three main compartments (black: presynaptic compartment ; dark gray: central compartment ; gray: postsynaptic compartment) are accessible though fluidic inlets / outlets and linked through structured axon valves microchannels (white and red). The presynaptic and postsynaptic compartments are seeded with neurons while the central compartment improves the fluidic isolation between the culture compartments and permits the examination of isolated axonal tracts. The white arrow indicates the imposed direction of axonal growth in the microchannels. The height of culture chambers is 50 μm while microchannels are 3 μm high. **(b)** Axon valves microchannels design and principle of unidirectional axonal filtration. In green, neurons originating from the presynaptic chamber can cross the microchannels array while in red, neurons originating from the postsynaptic chamber are either filtered by axonal traps or by arches microchannels. **(c)** Schematic representation of the experimental design. Hip^mTmG +/-^>Hip^mTmG -/-^ networks permitted to evaluate forward axonal growth (in green) while Hip^mTmG -/-^>Hip^mTmG +/-^ permitted to observe backward axonal growth (in red). **(d**,**e)** Representative epifluorescence microscopy fields of chimeric reconstructed neural networks after 19 days of culture. **(f)** Quantification of forward and backward axonal outgrowth in chimeric Hip>Hip networks, with individual points representing individual culture devices. The number n of individual culture devices is indicated on the graph, from N=3 individual experiments. Error bars show standard deviation.

While most studies assessing axonal filtration in asymmetric microchannels have evaluated axonal outgrowth when only one compartment is seeded, axons can serve as guides to neuronal processes growing in the opposite direction. We thus evaluated axonal filtration in chimeric networks, where one compartment was seeded with hippocampal primary neurons expressing the fluorescent protein mtdTomato (Hip^mTmG +/-^) neurons, and the other with non-fluorescent Hip^mTmG -/-^ neurons (Fig 1c). This allowed us to selectively observe forward or backward axonal outgrowth (Fig 1d and 1e). Importantly, we found that axon valves totally prevented backward axonal outgrowth while allowing forward axonal outgrowth and robust invasion of the postsyn compartment (Fig 1f, Supp Fig 2).

### Active anterograde trans-neuronal transfer of exogenous aSyn aggregates in reconstructed neural networks decreases with maturation

aSyn aggregates are taken up by neurons and transported anterogradely and retrogradely in their axons [6, 11, 14]. We used our novel experimental setup to quantitatively demonstrate synapses-dependent neuron to neuron transfer of aggregated aSyn from donor to recipient naïve neurons and subsequent seeding in the latter. Two distinct aSyn aggregates that differ in size and structural and biochemical characteristics [4], that spread with different efficiencies in vivo, [40] were generated: Oligomers (hOlig) and Fibrils (hFib) [35]. We followed the fate of the exogenous aggregates thanks to covalently bound fluorophores.

We first evaluated if exogenous aggregates transfer occurred from presyn to postsyn neurons, and if it was mediated by axonal transport. For that, we used two types of networks: either both compartments were seeded with Hip neurons (Hip>Hip), or only the postsyn compartment was seeded (Ø>Hip). We introduced 500 nM of hFib or hOlig (concentration of aSyn monomers) in the presyn compartment for 24 h starting at day in vitro (DIV)14, and monitored the presence of exogenous aggregates in postsyn neurons 3 days later, at DIV17 (Supp Fig 3a). Autofluorescence was observed in control untreated networks in the form of somatic puncta, which could not readily be visually distinguished from weak aggregates-associated fluorescence in postsyn neurons of treated networks (Supp Fig 3a). We thus systematically normalized our results with the fluorescent signal observed in untreated networks. For both types of aggregates, we observed a moderate but statistically significant ∼10 % increase in postsyn neurons fluorescence in treated versus untreated Hip>Hip networks (Fig 2d, “postsyn” graph). No increase in postsyn fluorescence was observed in treated Ø>Hip networks (Supp Fig 3b and c). This demonstrates that hFib and hOlig are actively and anterogradely transferred from neuron to neuron.

**Figure 2.**
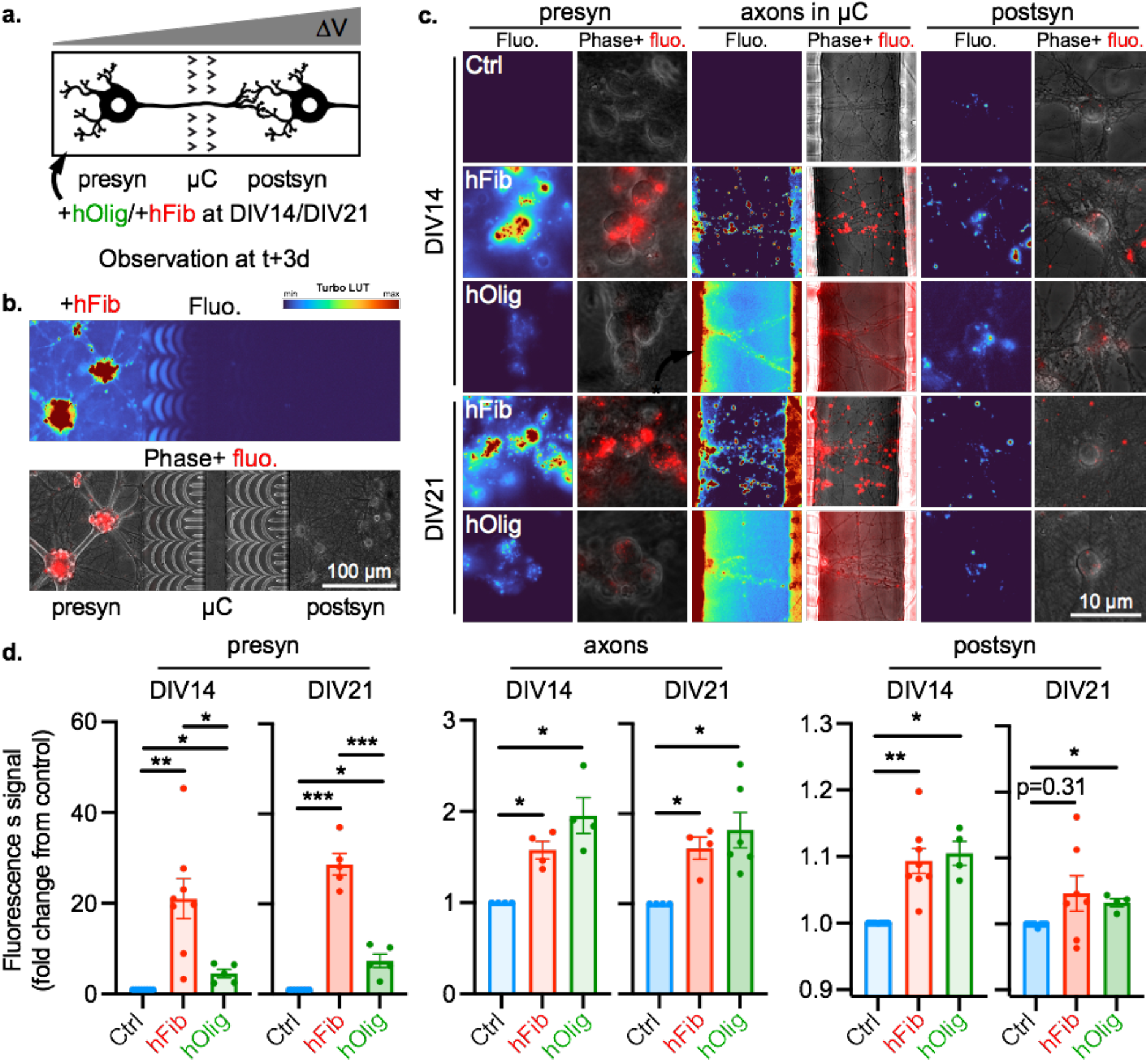
Trans-neuronal transfer of exogenous aggregates in immature and mature neural networks. **(a)** Schematic representation of the experimental design. The presynaptic compartment of Hip>Hip networks was spiked at DIV14 or DIV21 with control solution, 500 nM hFib or 500 nM hOlig, and aggregates associated fluorescence was monitored in neuronal somas in the postsyn and presyn compartments, and in axons in the central compartment 3 days later. **(b)** Low magnification epifluorescence microscopy field of a representative Hip>Hip network at DIV17, 3 days after presynaptic exposure to 500 nM hFib. Top, hFib and hOlig associated fluorescence is shown with the Turbo look-up table (LUT). Bottom, hFib and hOlig fluorescence is in red and Phase signal in gray. **(c)** Representative epifluorescence microscopy fields of presynaptic somas, axons in the central compartment and postsynaptic somas, 3 days after exposure of the presynaptic compartment to the indicated treatment. For each column, the left column highlights hFib associated fluorescence with the Turbo colormap, and the right column shows hFib fluorescence in red and Phase signal in gray. **(d)** Quantification of the aggregates associated fluorescence signal in neuronal somas in the presynaptic and postsynaptic compartments, and in axons in the central compartment. DIV14 presyn : n=24-33, N=5-8. DIV21 presyn : n=22-26, N=5-7. DIV14 axons : n=15-19, N=4. DIV21 axons : n=14-19, N=4-6. DIV14 postsyn : n=16-54, N=4-9. DIV21 postsyn : n=22-42, N=4-8. (n individual culture devices from N individual experiments). Brown-Forsythe test followed by Dunnett’s T3 multiple comparisons test. *** : p<0.001 ; ** : p<0.01 ; * : p<0.5. Error bars show standard error of the mean.

Despite the absence of data conclusively demonstrating the role of synaptic structures, the propagation of aSyn aggregation in neural networks is often considered “trans-synaptic” [10, 43]. Thus, the role of synaptic structures in the transfer of aSyn aggregates remains uncertain. Indeed, aSyn assemblies can be secreted in the extracellular medium by axonal projections [6]and traffic between neuron in immature neural networks, which are not synaptically connected [11]. We thus sought to address the impact of synaptic maturity on aSyn aggregates propagation. In our culture system, axonal projections from presyn neurons progressively invade the postsyn compartment, and trans-compartment synaptic connectivity increases between DIV14 and 21 in Hip>Hip networks (Supp Fig 4), in the same fashion as in cortico-striatal networks [24]. We hypothesized that if aSyn aggregates propagation is favored by the presence synaptic structures, we would observe a higher dissemination rate in more mature and synaptically connected Hip>Hip networks.

To test this hypothesis, we exposed the presyn compartment of DIV14 or DIV21 Hip>Hip networks to 500 nM of hFib or hOlig for 24h and monitored the distribution of aSyn aggregates 3 days after the initial introduction of aggregates in the system, 2 days after their removal from the medium (Fig 2a). The fluorescence of the media containing similar concentrations of hFib and hOlig differed by less than 2 fold (data not shown). While the spatial distribution of fluorescence remained strongly skewed toward the presyn compartment (Fig 2b), we observed fluorescence in axons and in postsyn neurons (Fig 2c). We previously showed that colocalization of hFib with neuronal somas following exposure through the culture medium was due to endocytosis, and will term this colocalization “direct uptake” in the rest of this manuscript [7]. Direct exposure of neurons resulted in a 21 fold increase of fluorescence in neuronal somas when spiking was performed at DIV14 and a 29 fold increase of fluorescence when it was performed at DIV21. hOlig direct uptake was significantly less efficient, with a 4.5 fold increase in fluorescence at DIV14 and a 7.3 fold increase at DIV21 (Fig 2d). This is in good agreement with previous results from cultured neurons showing that hFib bind much better to neuronal membranes than hOlig [47]. Aggregates associated fluorescence was also observed in axons in the central compartment, with a 60 % increase in signal in hFib treated networks at both DIV14 and DIV21, and a 95 % and 80 % increase in signal in hOlig treated network at DIV14 and DIV21, respectively (Fig 2d). While hFib fluorescence was mostly contained within axonal projections in this compartment, fluorescence outside axons was observed in hOlig treated networks (Fig 2c). This could be due to their passive diffusion to the central chamber. Indeed, while we showed that aSyn aggregates did not passively diffuse from the presyn compartment to postsyn neurons when the presyn compartment did not contain neurons (Supp Fig 3), some passive diffusion could occur between the presyn and axonal compartments. This might in turn impact our quantification of the anterograde transfer of aggregates from somas to axonal tracts. We next quantified the amount of trans-neuronal transfer. In DIV14 networks, both hFib and hOlig treatment resulted in a statistically significant 10 % increase in signal in postsyn neuronal somas. In DIV21 networks, despite a higher aggregates fluorescence in presyn somas, the signal increased more modestly in postsyn somas, 4.6 % and 3.2 % upon hFib or hOlig treatment, respectively (Fig 2d). We then asked if the presyn and postsyn increase in global neuronal fluorescence was carried by a sub-population of “super-uptakers”. We plotted the distribution of hFib and hOlig fluorescence in individual somas and found it followed a unimodal distribution in presyn and postsyn neurons, both at DIV14 and DIV21 (Supp Fig 5). This argues against the existence of neuronal populations that take up with different efficiencies hFib or hOlig.

Because the quantity of signal associated with presyn neurons differed greatly between hFib and hOlig exposed networks, we expressed our result as “Transmitted Excess Signal” (TES): a metric which expresses the fluorescent aggregates signal in the axonal or postsyn compartments as a percentage of the signal detected in the presyn or axonal compartment (see Methods for details). We plotted TES for hFib and hOlig as a function of culture maturity (Fig 3a). This metric allows to specifically assess the efficiency with which aggregates associated with presyn neurons are then transported in axons and transferred to postsyn neurons, independently of the efficiency of the rate-limiting steps of fixation and endocytosis in presynaptic somas (Fig 3a, Fig 3c). The hOlig TES from presyn to axons and presyn to postsyn are higher than for hFib, both at DIV14 and DIV21 (Fig 3b). This suggests that while hOlig are taken up to a lesser extent than hFib, their axonal transport and their transfer from soma to soma are more efficient. The axons to postsyn TES is similar for both hOlig and hFib, which suggests that the two kinds of aggregates transfer from axonal projections to secondary neurons with similar afficiency. Additionally, while these differences were not statistically significant, we noted that the axons to postsyn TES values for both aggregates are lower at DIV21 compared to DIV14, suggesting that network maturity and synaptic maturation did not increase transfer from axonal projections to secondary neurons. We also noted that the presyn to axons TES of hOlig decreased with network maturity.

**Figure 3.**
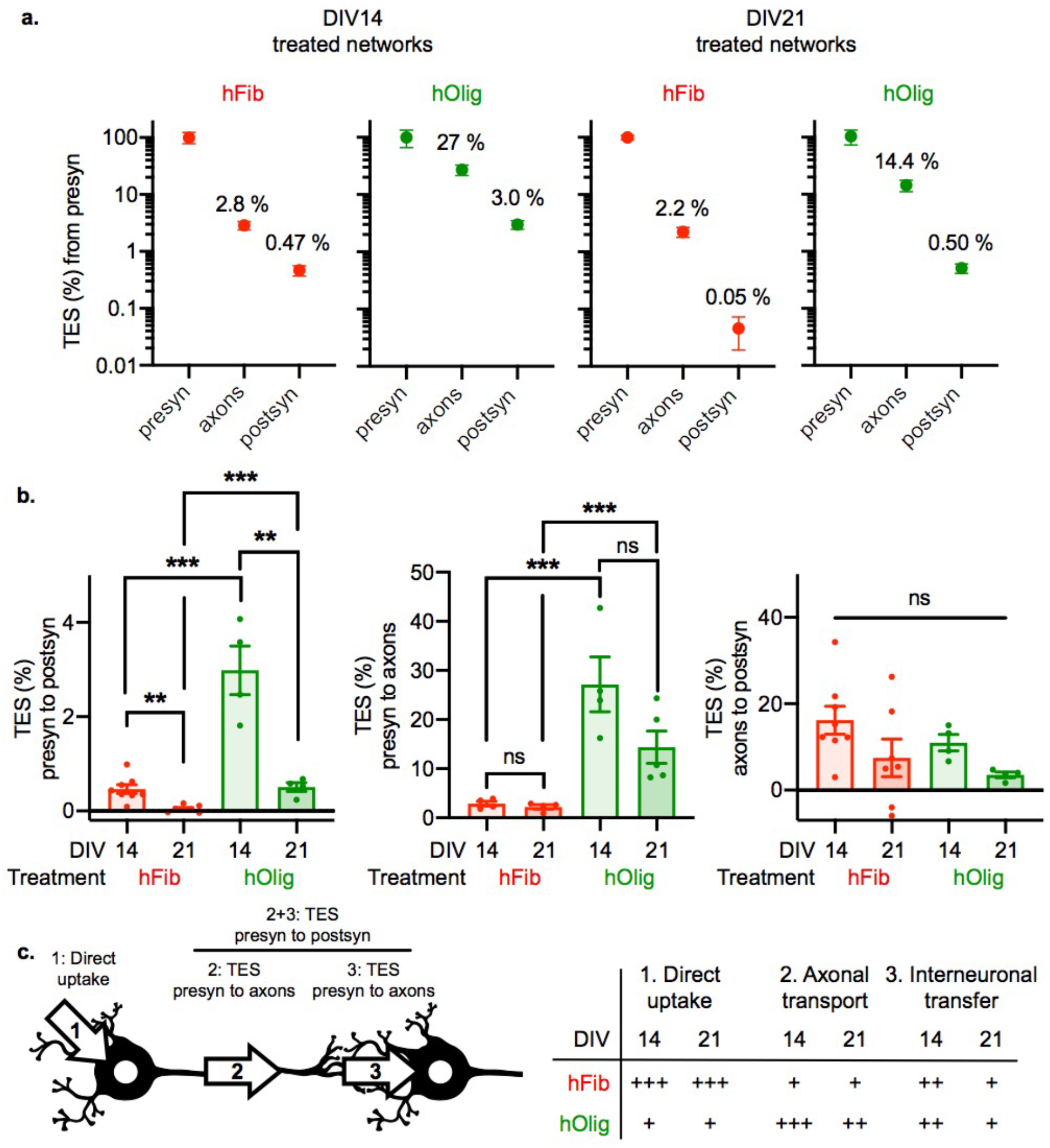
A small fraction of presynaptic aggregates are transferred to postsynaptic neurons, and network maturity decreases trans-neuronal transfer of exogenous aggregates. **(a)** Transmitted Excess Signal (TES) profiles generated from results reported in Fig 4 (see Methods for calculation). Mean and standard error of the mean are shown in log scale. **(b)** TES in between each compartment. Two way ANOVA followed by Tukey’s multiple comparisons test.. *** : p<0.001 ; ** : p<0.01 ; * : p<0.5. **(c)** Summary of observed transmission rates in between each compartments. Left, scheme of the different steps of an axon dependent trans-neuronal transfer of exogenous aggregates. Right, summary of the efficiency (from + : inefficient to +++ : efficient) of the different steps depending on the nature of the exogenous aggregate and the maturity of the network. Step 1 refers to the two rightmost graphs in Fig 4d. Error bars show standard error of the mean.

In conclusion, these data support that following fixation to presynaptic somas, hOlig are more efficiently transported to postsynaptic neurons than hFib. This is to some extent expected and in good accordance with previous observations made *in vivo* [40]. Interestingly, this difference in transfer efficiency appears to result from a higher rate of transfer from neuronal bodies to axons (Fig 3b), while the TES from axons to secondary neurons does not significatively differ in between the two aggregates. Thus, higher network maturity and synaptic connectivity decreased the transfer of both polymorphs from presyn to postsyn, through the combined effect of reduced axonal transport and reduced axon to neuron transfer (Fig 3c).

### Anterograde trans-neuronal propagation of aSyn seeding is inefficient and starts in the axons of postsynaptic neurons

Amplification of pathological aggregates through seeding of soluble aSyn is thought to be the motor of the long term and long-distance propagation of aggregation in brain networks. However, neural network topology alone does not predict the path of aggregates propagation in brain networks, and the efficiency of retrograde versus anterograde aggregates spread remains elusive [50]. We thus sought to evaluate the efficiency of trans-neuronal aSyn aggregation propagation in our setup uniquely suited for studying pure anterograde propagation of protein aggregates with prion-like properties.

We previously showed selective phosphorylation of Ser 129 residue upon seeded aggregation of endogenous aSyn [7, 12]. To assess in a quantitative manner seeding of endogenous aSyn by exogenous aSyn aggregates, the presyn compartment of DIV14 Hip>Hip networks was exposed to 500 nM hFib or hOlig for 24 h, fixed and stained by anti-S129 phosphorylated aSyn (pSyn) antibody 10 days later (Fig 4a). Exposure to hFib, but not hOlig led to the formation of numerous pSyn positive inclusions in all compartments (Fig 4b, c and d). Propagation of aggregation was dependent on the presence of neurons in the presyn compartment, supporting that seeding species do not diffuse passively in between compartments in the 10 days following initial spiking, given that a pressure gradient is maintained in the microfluidic system (Supp Fig 6). It was previously reported local spiking of bidirectional compartmentalized neural networks led to the formation of abundant somatic aggregates in secondary neurons in a few days in a triple compartmentalized culture setup [52]. In over 20 hFib treated Hip>Hip networks carefully examined by epifluorescence and confocal microscopy, we observed no somatic inclusions in postsyn neurons. In contrast, introduction of hFib on one side of bidirectional compartmentalized networks led to the accumulation of somatic pSyn inclusions in 1 to 3 second order neurons per network, in n=10 networks from N=2 separate experiments (Fig 5). This suggests that previously reported trans-neuronal propagation of aggregation in triple compartmentalized networks might either result from a higher efficiency of retrograde trans-neuronal propagation, or, as observed here, from the presence of axonal projections in between the two most peripheral compartments. While we observed no somatic inclusions in postsyn neurons, we hypothesized that inclusions might first form in neuritic projections, as neuropathological observations suggest [54]. Indeed, the subcellular localization of pSyn rich inclusions in the postsyn compartment of Hip>Hip networks was ambiguous. Confocal imaging revealed that pSyn inclusions colocalized with axons. We estimated that colocalization of pSyn with MAP2 could not be attributed with certainty to seeding in the secondary neurons, and might instead be due to the fasciculation of axonal and dendritic processes (Fig 6a, b, c and d). In order to determine if some of the axonal inclusions that we detected in the postsyn compartment were located in axons from the postsyn neurons, we selectively infected the postsyn population with an adeno-associated viral vector encoding the enhanced green fluorescent protein (AAV-GFP) (Fig 6e). While the interpretation of most GFP-pSyn colocalizations were made ambiguous by the fasciculation of numerous axonal projections from both presyn and postsyn axons, we detected the unambiguous colocalization of pSyn inclusions with isolated GFP positive axons in 1 out of 4 hFib treated networks used in this experiment (Fig 6f and g). These data support that anterograde transfer of aSyn seeds from presynaptic neurons leads to trans-neuronal aggregation starting in the axons of postsynaptic neurons.

**Figure 4.**
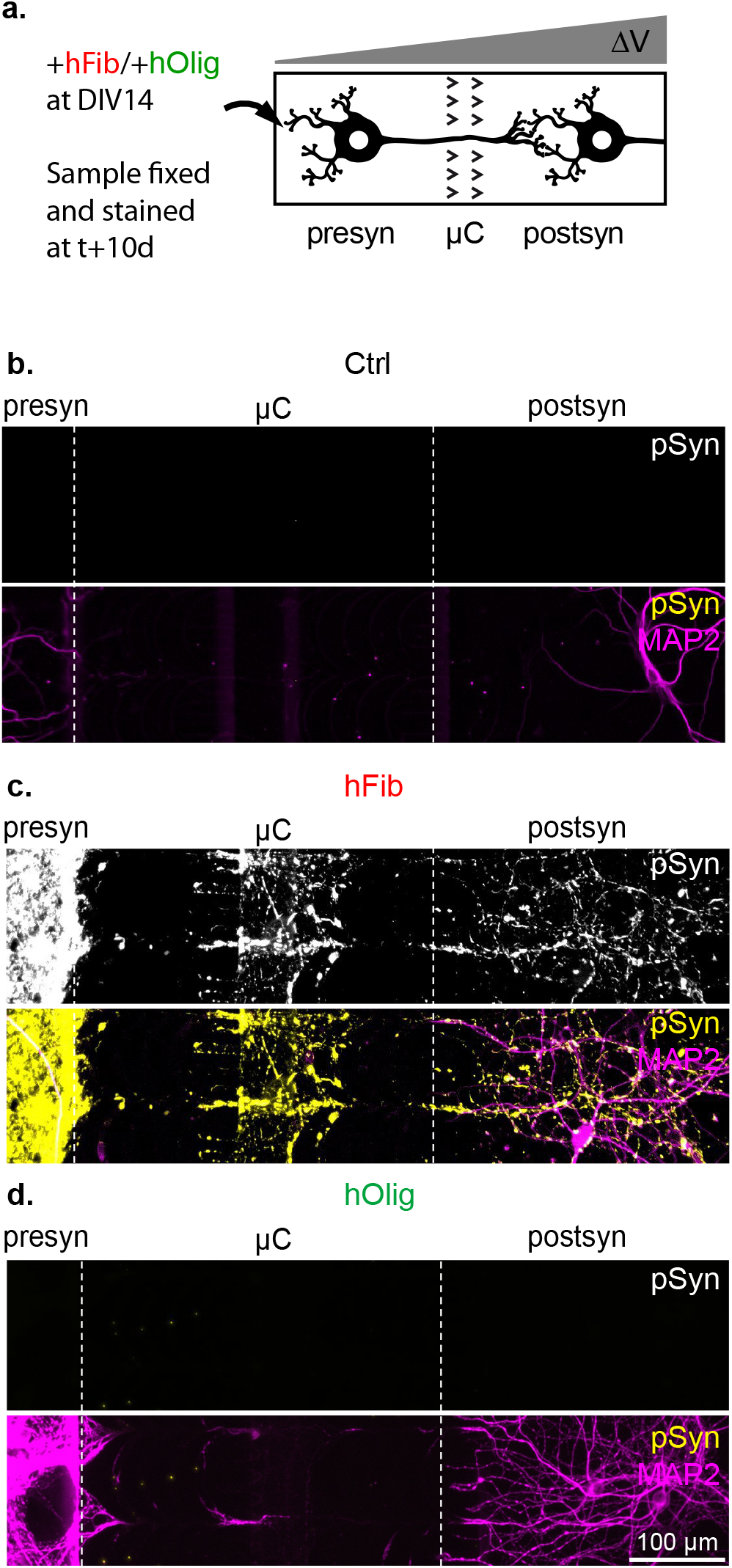
Propagation of endogenous aSyn aggregation after exposure of the presynaptic compartment of oriented networks to aSyn Fibrils. **(a)** Schematic representation of the experimental design. The presynaptic compartment of Hip>Hip networks was spiked at DIV14 with control solution, 500 nM hFib or 500 nM hOlig, and cultures were fixed and stained 10 days later. **(b**,**c**,**d)** Representative epifluorescence fields of Hip>Hip networks after presynaptic exposure to **(b)** control treatment, **(c)** hFib, **(d)** hOlig.

**Figure 5.**
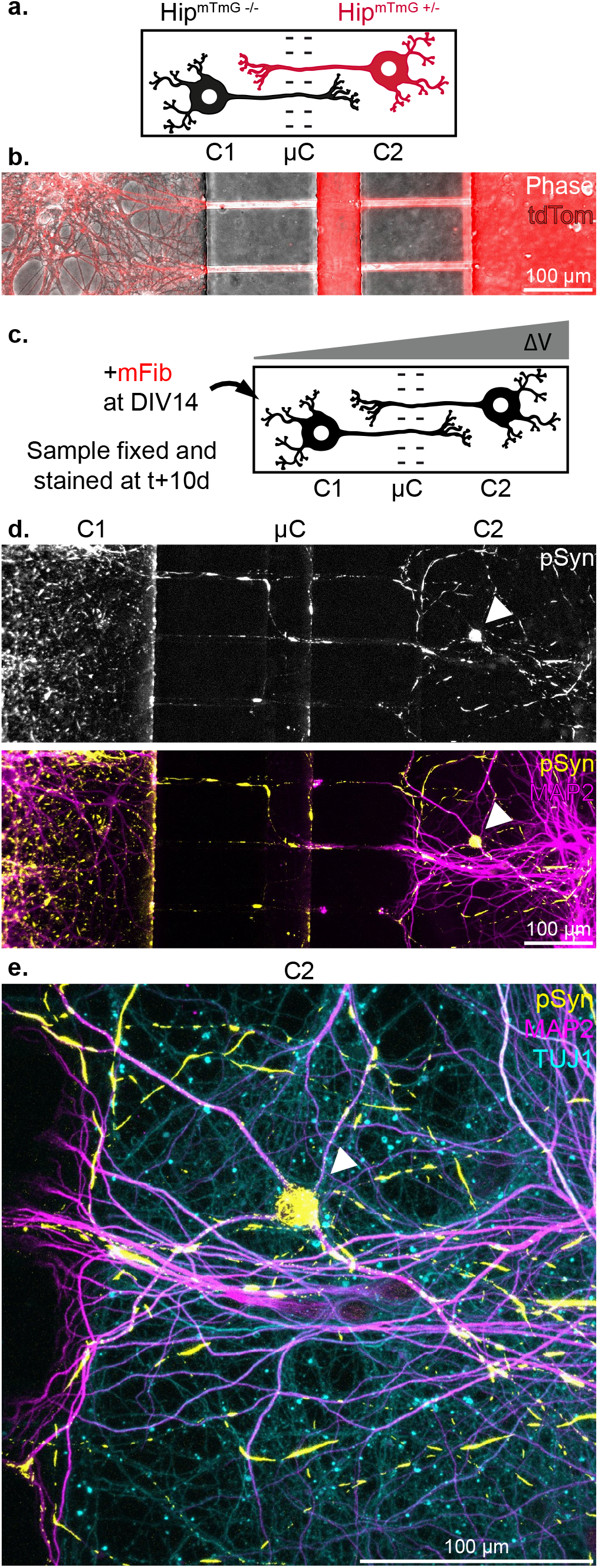
Retrograde axonal projections permit consequent aSyn aggregation propagation. **(a)** Schematic depiction of a Hip^mTmG -/-^-Hip^mTmG +/-^ bidirectional network. Straight 10 μm wide microchannels, in a device otherwise similar to the UND in other aspects, allow axons to grow in both directions. **(b)** Representative image of axonal tracts connecting the two culture compartments in a Hip^mTmG -/-^-Hip^mTmG +/-^ bidirectional network. **(c)** Schematic representation the experimental design. The compartment 1 (C1) of a Hip-Hip network was spiked at DIV14 with 500 nM of mFib. A pressure gradient was maintained during the whole culture after spiking to prevent passive diffusion to the second compartment. The culture was fixed 10 days later. **(d)** Representative image of a Hip-Hip network treated as explained in (c). The white arrow points to a C2 neuron containing numerous somatic pSyn inclusions. **(e)** Confocal imaging of the neuron highlighted in (d), highlighting the unambiguous somato-dendritic accumulation of pSyn inclusions.

**Figure 6.**
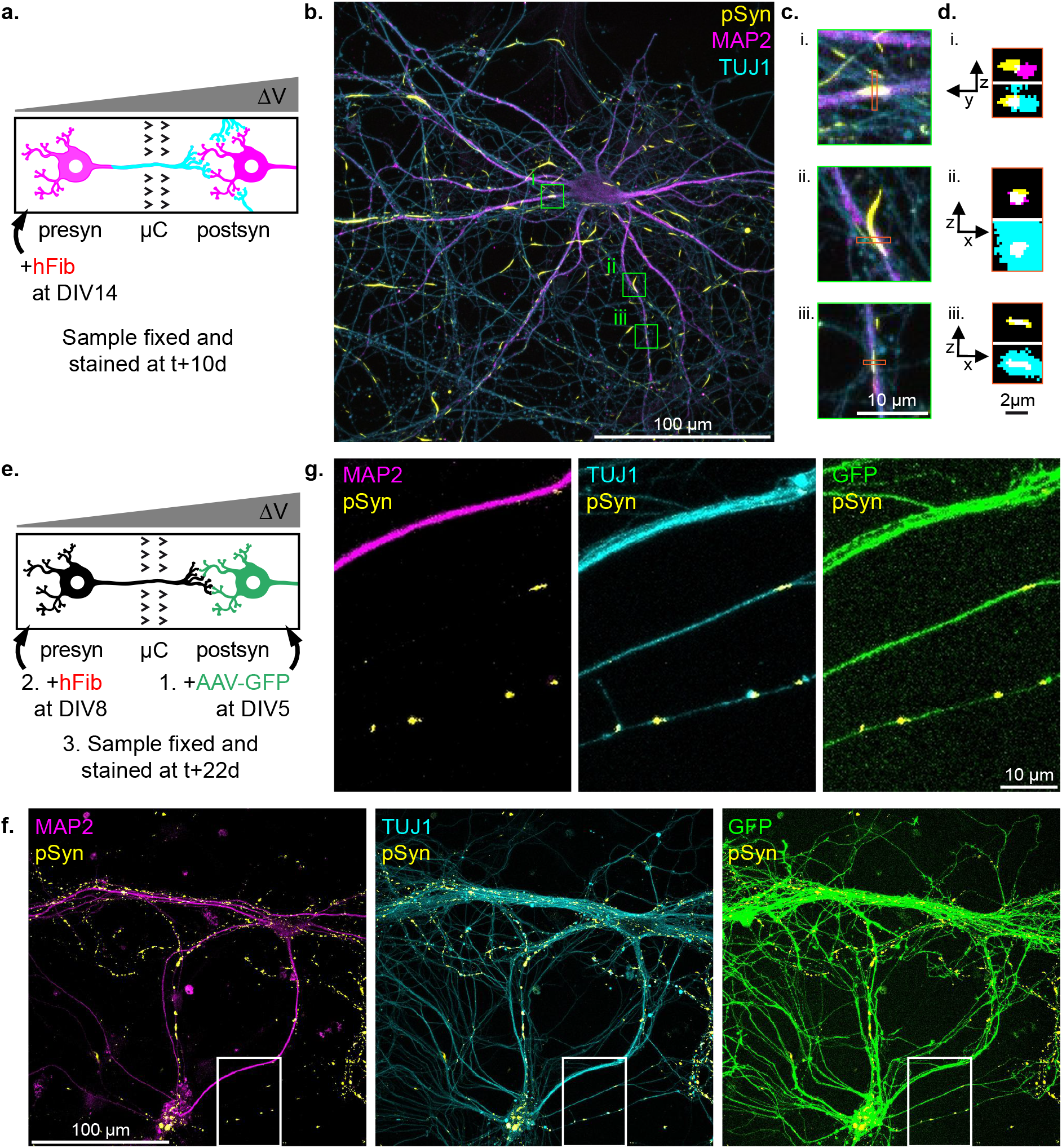
Exposure of the presynaptic compartment of reconstructed networks to aSyn Fibrils leads to modest and purely axonal aSyn aggregation in postsynaptic neurons. **(a)** Schematic representation of the first experimental design. The presynaptic compartment of Hip>Hip networks was spiked at DIV14 with 500 nM hFib, and cultures were fixed and stained 10 days later. **(b)** Representative confocal microscopy field of a postsynaptic neurons 10 days after hFib spiking of the presynaptic compartment. pSyn inclusions localization is ambiguous and appears mostly axonal. **(c)** Higher magnification view of the fields highlighted in (b), showing on pSyn inclusions that appear totally (i.,iii.) or partially (ii.) colocalized with MAP2. **(d)** (x,z) or (y,z) view of the regions highlighted in (c). It appears that pSyn inclusions which do not appear to be contained in neuritic projections (ii.) colocalizes as much or more with the MAP2 signal than inclusions which appear to be contained in MAP2 from the (x,y) view. **(e)** Schematic representation of the second experimental design. To determine if some of the pSyn inclusions are located in the axons of postsynaptic neurons, the postsynaptic neurons of Hip>Hip networks were exposed to AAV-GFP at DIV3, days before axons reach from the presynaptic to the postsynaptic compartment. The networks were then processed as previously.**(f)** One of the rare confocal microscopy fields in which an unambiguous colocalization of GFP and pSyn signals was observed. **(g)** Magnification of the region from fields in (f) containing the unambiguous co-localization of GFP, pSyn and TUJ1 signals in isolated axonal projections.

### Presynaptic neuronal subtypes do not affect exogenous aggregates transfer, but determine the magnitude of aSyn seeding

aSyn aggregation spreads within the brain of patients in distinct neural networks, and some have suggested that the characteristics of neuronal populations making up the networks impact the efficiency of rate-limiting steps in this propagation, independently of the synaptic strength between those populations [20, 21, 50]. Supporting this notion, we previously established a relationship between the seeding propensity of exogenous murine aSyn Fibrils (mFib) and endogenous aSyn expression level using distinct neuronal cultures (hippocampal, cortical and striatal neurons) [7]. To evaluate if the nature of the neuronal population also impacts exogenous aSyn seeds propagation, we constructed Hip>Hip, Hip^SNCA -/-^>Hip^SNCA +/+^ and cortico-hippocampal (Cx>Hip) networks (Fig 7a). We treated the presyn compartment of these networks at DIV14 with 500 nM mFib and followed aggregates transfer to postsyn neurons 1 day after treatment. We observed mFib associated fluorescence in postsyn neurons in the 3 types of networks we constructed (Fig 7b). The increase in fluorescence from background signal was of 8 %, 7 % and 4 % in Hip>Hip, Hip^SNCA -/-^>Hip^SNCA +/+^ and Cx>Hip, respectively. No statistically significant differences were observed when individual culture devices instead of experimental means were considered in the statistical analysis (Fig 7c). We next assessed pSyn accumulation 10 days after treatment, and found, as previously published [7], that pSyn signal was absent in Hip^SNCA -/-^, low in Cx, and high in Hip neurons, with the quantity of pSyn in the postsyn compartment reflecting the quantity of pSyn detected in presyn (Fig 7d, e, f). We did not observe any somatic pSyn inclusions in postsyn neurons in any of the networks considered. Careful examination of 9 Hip^SNCA -/-^>Hip^SNCA +/+^ networks, did not reveal any trace of pSyn in postsyn axons and dendrites.

**Figure 7.**
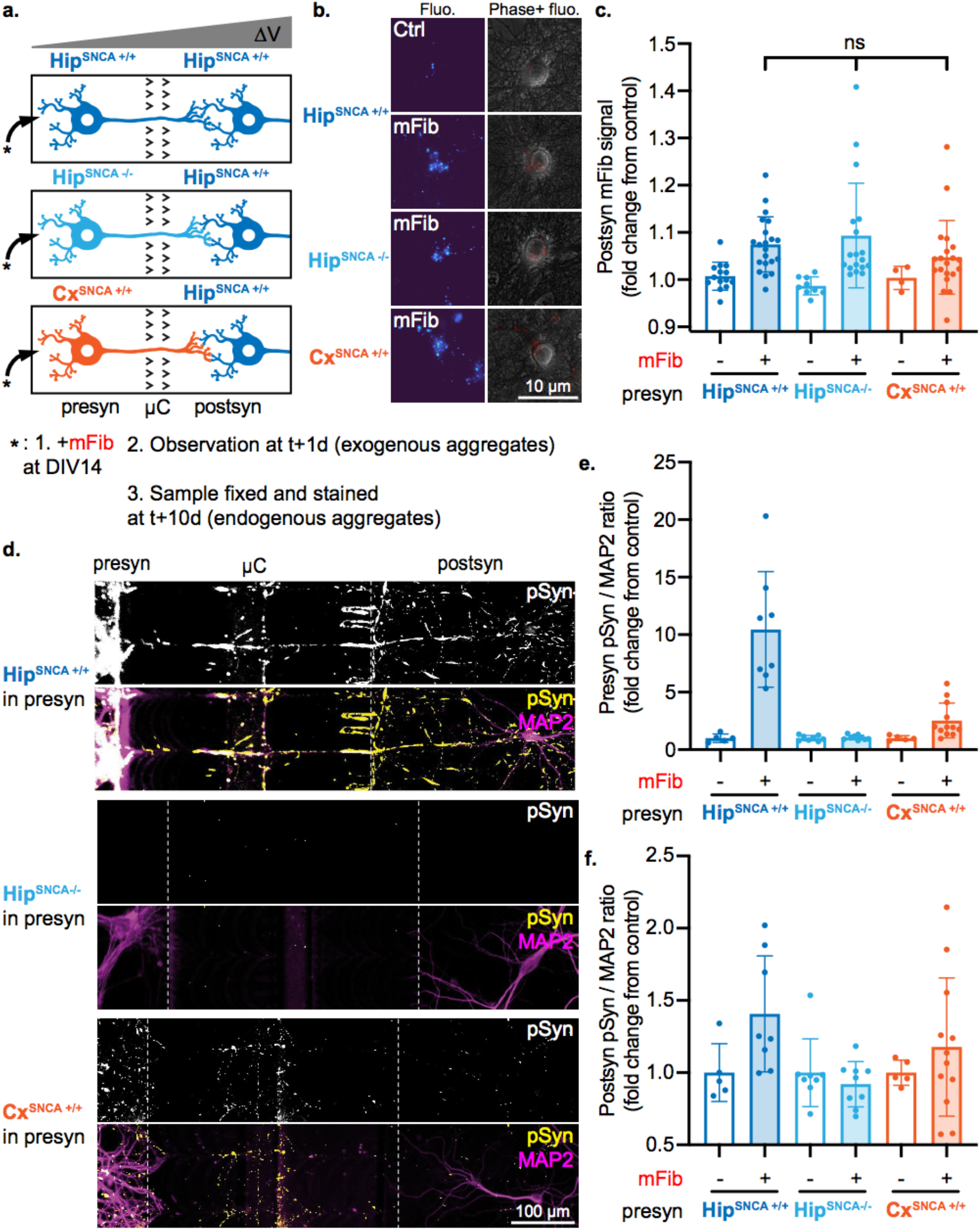
Presynaptic identity does not impact fast aggregates transfer, but does impact endogenous aSyn aggregates seeding. **(a)** Schematic representation of the experimental design. The presynaptic compartment of Hip^SNCA +/+^>Hip^SNCA+/+^, Hip^SNCA -/-^>Hip^SNCA+/+^ and Cx^SNCA +/+^>Hip^SNCA+/+^ networks was spiked at DIV14 with 500 nM mFib, aggregates associated fluorescence was monitored in neuronal somas in the postsyn compartment 3 days after treatment, and networks were fixed and stained 10 days after treatment. **(b)** Representative epifluorescence microscopy fields of postsynaptic neurons 1 day after presynaptic exposure to mFib. Left, mFib associated fluorescence is shown with the Turbo colormap. Right, mFib fluorescence is in red and Phase signal in gray. **(c)** Quantification of mFib associated fluorescence in postsynaptic neurons. Results from individual culture devices are shown as individual data points. Kruskall-Wallis test followed by Dunn’s multiple comparisons test was performed selectively on data from mFib treated networks. n=18-21 devices from N=3 individual experiments in mFib treated networks, and n=4 to 14 devices from N=3 individual experiments in ctrl networks. **(d)** Representative epifluorescence microscopy fields of stained networks, 10 days after presynaptic exposure to mFib. **(e**,**f)** Quantification of the pSyn/MAP2 signals ratio in **(e)** presynaptic and **(f)** postsynaptic compartments. n=5-12 culture devices from N=1-2 individual experiments. Error bars show standard deviation.

We conclude from these results that Cx and Hip neurons transfer exogenous aSyn fibrils to postsynaptic neurons at similar rates. We further conclude that differential aggregation of endogenous aSyn within wild type Cx and Hip neurons does not affect seeding in postsynaptic neurons. Exogenous aSyn aggregates propagation propensity of in aSyn knockout mice overexpressing locally human aSyn has been reported to be higher than in WT mice [19]. Our results suggest that differential, yet physiological, aSyn expression does not impact trans-neuronal transfer of exogenous aggregates.

## Discussion

Pathological aSyn spreading is widely considered “trans-synaptic” given that regions that sequentially develop LP are most often synaptically connected and that neuroanatomical connectivity supports the spread of LP better than a nearest neighbor scheme [5, 30]. aSyn assemblies have been shown to traffic between neuronal cells after release in the culture medium naked or within exosomes [36], through TNT [1, 42], and possibly immature synapses [11]. Thus, while the cellular and molecular mechanisms underlying inter-neuronal transfer remains unclear, quantifying the efficiency of seeds propagation between synaptically connected neurons is important for both accurately modeling the spread of aSyn aggregates and pinpointing biomolecular cues involved in that transfer. Here, using a newly developed microfluidic device allowing the reconstruction of fully oriented binary neuronal networks with virtually no retrograde connections, we demonstrate that the prion-like dissemination of aSyn aggregates between two synaptically interconnected neurons is a slow process, highly dependent on the size of aSyn aggregates.

### Quantification of aSyn aggregates spread

Our results provide a first quantitative estimate of exogenous aSyn aggregates anterograde transfer between 2 interconnected neurons. While aSyn seeds are efficiently taken up by neurons, most exogenous aggregates remain within the somato-dendritic compartment, with only ∼ 2% of the Fibrils found within axonal shafts of presynaptic neurons. While 0.5 % of presyn hFib was detected in postsynaptic neurons 3 days after presynaptic inoculation in DIV14 networks, hOlig spread much more efficiently than hFib between neurons, with 3 % fluorescence transfer from presynaptic to postsynaptic neurons. Assuming a linear correlation between fluorescence increase and aggregates concentration, we estimate that in the 3 days following exposure of the presynaptic compartment of Hip>Hip networks to 500 nM of hFib, the postsynaptic compartment was exposed to 2.5 nM and 250 pM of hFib, in DIV14 and DIV21 networks, respectively. With an average of ∼100 axons innervating the postsyn compartment, a single neuron whose somatodendritic compartment is exposed to 500 nM of hFib anterogradely transfers ∼8 pM of hFib per day during this period. This rate falls to ∼0.8 pM of hFib per day in DIV21 networks. Following the same reasoning, the rate of anterograde hOlig transfer per axon is of ∼50 pM/day in DIV14 networks and ∼8 pM/day in DIV21 networks in the 3 days following exposure of the presynaptic compartment to 500 nM of the aggregate

Surprisingly, the transfer of both hFib and hOlig was lower in DIV21 networks compared to DIV14 networks, despite the fact that synaptic connectivity between the two populations increased with time of culture. This suggests that intact synaptic structures are less permeable to large aSyn aggregates as compared to smaller ones and impede inter-neuronal transfer. This is in agreement with previous observations made in vivo showing that aSyn Fibrils spread to second order neurons to a much lesser extent than monomeric or aSyn Oligomers after delivery within the olfactory bulb [38, 40]. Importantly, in our paradigm, neither the synapse identity (cortex vs hippocampal) nor the genetic status (WT vs aSyn knockout) of presynaptic structures significantly modified trans-neuronal spreading efficiency.

### Efficiency of trans-neuronal seeding

Previous works aiming at investigating aSyn aggregates prion-like propagation within in vitro reconstruction neural networks suffered from several limitations, such as an absence of synaptic connectivity [11], or uncertainty as to the direction of axonal growth in the first generation of microfluidic devices [29, 52]. The results we report here using a novel microfluidic culture system to generate synaptically mature and fully unidirectional neural networks, in which the trans-neuronal transfer of aSyn aggregates can unequivocally be attributed to active anterograde axonal transport are in sharp contrast with reports of efficient trans-neuronal spreading in networks reconstructed with conventional non-filtering microfluidic devices [29, 52]. Some of those results have since then been contradicted [9]. This difference in propagation efficiency might arise from direct exposure of axons originating from neurons in the distal chamber of the microfluidic device that invade the proximal chamber, where aSyn aggregates are added. This is strongly suggested by our results showing significant endogenous aSyn seeding in secondary neurons grown in microfluidic devices with straight micro-channels. While this calls for attention upon use of microfluidic systems, this also evidences direct uptake of aSyn by axonal termini and very efficient retrograde transfer of aSyn seeds towards neuronal cell bodies. A partial correlation between synaptic connectivity and the pattern of propagation of aSyn aggregation was recently evoked [20, 21, 50]. Our results suggest that direct axonal capture of injected aggregates may fully account for these observations.

However, many aspects of cultured neurons physiology other than synaptic connectivity evolve in the weeks following plating and might modify transfer efficiency. First, changes in the neuronal transcriptome might affect binding, uptake and clearance of exogenous aSyn aggregates. Second, immature neurons exhibit higher exocytic and endocytic activities, which might result in heightened uptake and secretion of exogenous aggregates, and could explain why transfer is more efficient in young networks [51]. Third, astrocytes proliferate over time in neuronal cultures and they have been shown to take up and degrade exogenous aSyn assemblies [26], thus, diverting aSyn assemblies transfer from neuron-neuron to neuron-astrocytes in older cultures. However, we observed that aggregates direct uptake was not diminished by culture maturation, arguing against a higher rate of aggregates capture by astrocytes with culture maturation (Fig 2).

### Overall pattern of spreading: temporal and spatial consideration, relation to the connectome spread

aSyn prion-like dissemination in brain networks theoretically encompasses 2 sequential steps involving 1) trans-neuronal spread of aSyn seeds and 2) amplification of seeds through a templated nucleation process. In previous studies we demonstrated a seeding efficiency dependence on seeds concentration and on endogenous aSyn expression in recipient cells [7, 46]. Direct exposure of neurons to hFib for 1 week led to the important accumulation of pSyn aggregates both at the somato-dendritic level and in the distal axonal shafts. Detecting pSyn aggregates in post synaptic neurons turned out to be difficult as most of pSyn structures in post synaptic chambers belonged to the presynaptic axons. Nevertheless, using viral vectors to trigger GFP expression in postsynaptic neurons, we were able to detect a small amount of endogenous aSyn inclusions in postsynaptic neurons. Strikingly, all inclusions we detected were located in the distal axons of post synaptic neurons with no sign of pSyn accumulation in their somato-dendritic compartments. These results are in line with previous data showing that, in vivo, LB accumulation starts in branched axons and follows a “centripetal” pattern towards the nucleus in neurons [21, 23, 43, 53]. These observations have been consistently reproduced in primary neuronal cultures [7, 57]. The limited amounts of pSyn in postsynaptic neurons axons might be the consequence of the amount of hFib transferred to the second order neurons, which we estimated earlier to be of 2.5 nM over a 3 days period in DIV14 networks.

The vulnerability to seeding of a given neuron in a brain network depends on the membrane proteins it expresses that bind seeds, endogenous aSyn expression level, the rate of seeds uptake and clearance by the recipient neuron [2, 7] but also, and most important, on the release of seeds from affected neighbor neurons. Overall, our results suggest that aSyn aggregates trans-neuronal spread depends on 2 distinct rate limiting steps encompassing 1) a size-dependent trans-neuronal traffic of aSyn aggregates and 2) a dependence on endogenous aSyn concentration leading to compartmentalized aSyn aggregation in distant areas of second order neurons. These aspects of aSyn aggregation might turns important for interpreting LB distribution in vivo and the apparent discrepancy between LB accumulation and the connectome. Indeed, if one was to count all (dendritic, somatic and axonal) inclusions in a given brain region to determine its vulnerability to PD, the presence of axonal afferents from affected neurons which somas are located in another region might lead to an overestimation of the vulnerability of the first region.

## Conclusion

By achieving for the first time the fully unidirectional filtration of axonal projections, we developed a new culture system for studying the anterograde propagation of aSyn aggregates in neural networks. We used this system to quantitatively study several aspects of aSyn aggregates propagation that are not easily accessible in conventional models, and obtained original data on elusive aspects of aSyn aggregates spreading. We notably determined that a very small fraction of aggregates made their way to secondary neurons. Surprisingly, we observed that more mature DIV21 networks with a higher synaptic connectivity showed a reduced capacity to transfer hFib and hOlig from neuron to neuron than immature DIV14 networks in the 3 days following exposure of the presynaptic compartment to aggregates. We also showed that the nature of the presynaptic population did not significantly modify the efficiency of seeds transfer. Finally, we obtained data supporting that anterograde trans-neuronal spreading of Fibrils leads to seeding of aggregation in postsynaptic neurons starting in their axonal projections. Overall, this work highlights the potential of tailor-made sophisticated culture system for deciphering new aspects of neuronal networks pathophysiology.

## Acknowledgements

J.C. was funded through a French Ministry of Research and Education doctoral fellowship and through the HoliFab European Commission project (Grant ID #760927). N.A.L. was funded through a “Fondation Recherche Alzheimer” doctoral fellowship. This work has also received the support of Institut Pierre-Gilles de Gennes (“Investissements d’avenir” program ANR-10-IDEX-0001-02 PSL, ANR-10-LABX-31 and ANR-10-EQPX-34) and of Région Ile-de-France (DIM Elicit), the European Union Joint Programme on Neurodegenerative Disease Research and Agence National de la Recherche (contracts PROTEST-70, ANR-17-JPND-0005-01 and Trans-PathND, ANR-17-JPND-0002-02) and the European Union’s Horizon 2020 research and innovation program and European Federation of Pharmaceutical Industries and Associations (EFPIA) Innovative Medicines Initiative 2 grant agreements No 116060 (IMPRiND) and No. 821522 (PD-MitoQUANT).

We thank Dr. Yassine GHOUZAM for adapting the Mask-RCNN method for detecting neuronal somas from Phase micrographs. We thank Dr. Benjamin LASSUS for helpful comments. We thank the members of the Axonal Regeneration and Growth team and of the Macromolecules and Microsystems in Biology and Medicine team for numerous discussions about the manuscript. We thank the Technical Core of IPGG (Institut Pierre-Gilles de Gennes), the Microscopy Core of IBPS (Institut de Biologie Paris-Seine) and the Electron Microscopy Facilities of I2BC (Institute for Integrative Biology of the Cell).

## Authors contributions

J.C. designed and fabricated microfluidic devices, generated primary neuronal cultures, performed synuclein seeding experiments, performed immunofluorescent staining, acquired epifluorescence and confocal images, analyzed the data, performed the statistical testing, generated the figures and wrote the first draft of the manuscript. N.A.L. fabricated microfluidic devices, generated primary neuronal cultures, performed viral transduction assays and synuclein seeding experiments, performed immunofluorescent staining and acquired confocal images. L.B. generated, characterized and labelled exogenous aSyn fibrillar assemblies. J.C., C.V., R.M. and J.M.P. participated in the conception of the experimental plan. J.C., C.V., R.M. and J.M.P. wrote the manuscript. All authors read and approved the final manuscript.

