## Supplemental Data for "Quantitative study of alpha-synuclein prion-like spreading in fully oriented reconstructed neural networks reveals non-synaptic dissemination of seeding aggregates"

**Title**

### SUPPLEMENTARY DATA

**Authors + Institutional addresses**

Josquin COURTE, PhD

Sorbonne Universités, Faculté des Sciences et Technologie, CNRS UMR 8246, INSERM U1130, Neurosciences Paris Seine, Institut de Biologie Paris Seine, Paris, 75005, France

Institut Curie, CNRS UMR 168, Université PSL, Sorbonne Universités, Paris 75005, France

The Eli and Edythe Broad CIRM Center, Department of Stem Cell Biology and Regenerative Medicine, University of Southern California Keck School of Medicine, Los Angeles 90033, USA

Ngoc Anh LE

Sorbonne Universités, Faculté des Sciences et Technologie, CNRS UMR 8246, INSERM U1130, Neurosciences Paris Seine, Institut de Biologie Paris Seine, Paris, 75005, France.

Luc BOUSSET, PhD

Institut François Jacob, (MIRCen), CEA and Laboratory of Neurodegenerative Diseases, CNRS, 92260 Fontenay-aux-Roses, France.

Ronald MELKI, PhD

Institut François Jacob, (MIRCen), CEA and Laboratory of Neurodegenerative Diseases, CNRS, 92260 Fontenay-aux-Roses, France.

Catherine VILLARD, PhD

Institut Curie, CNRS UMR 168, Université PSL, Sorbonne Universités, Paris 75005, France

Jean-Michel PEYRIN, PhD

Sorbonne Universités, Faculté des Sciences et Technologie, CNRS UMR 8246, INSERM U1130, Neurosciences Paris Seine, Institut de Biologie Paris Seine, Paris, 75005, France

**Corresponding author**

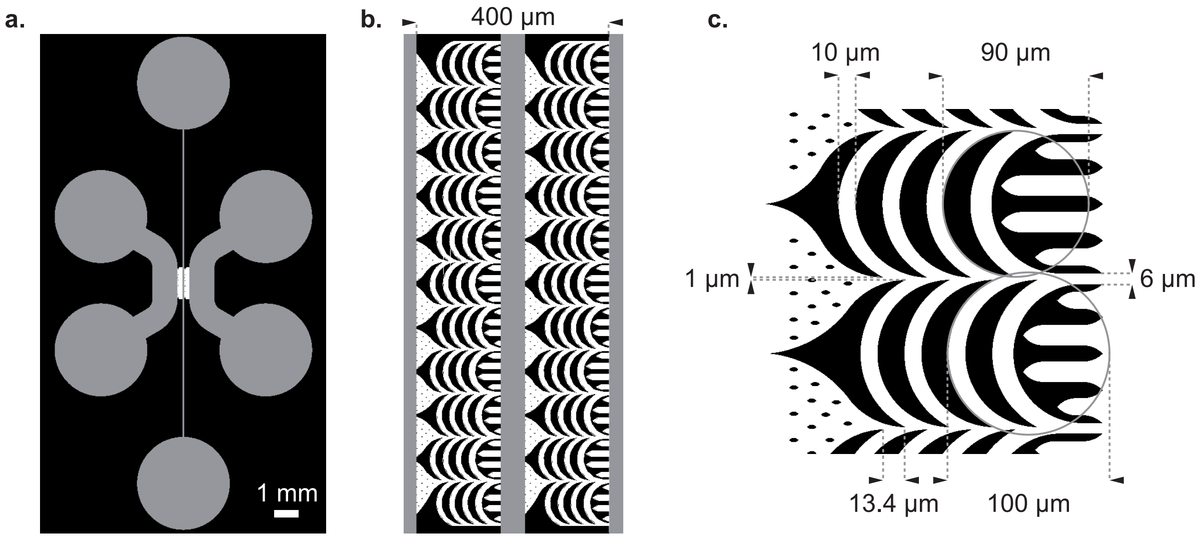


**Supplementary Figure 1**

**Dimensions of axonal filtration microchannels**

**(a)** Design of a single device for oriented network reconstruction. In grey: 50 µm high compartments, in white: 3 µm high microchannels. **(b)** Zoom on the microchannels. **(c)** Zoom on a single microchannel motif. Critical dimensions are highlighted.

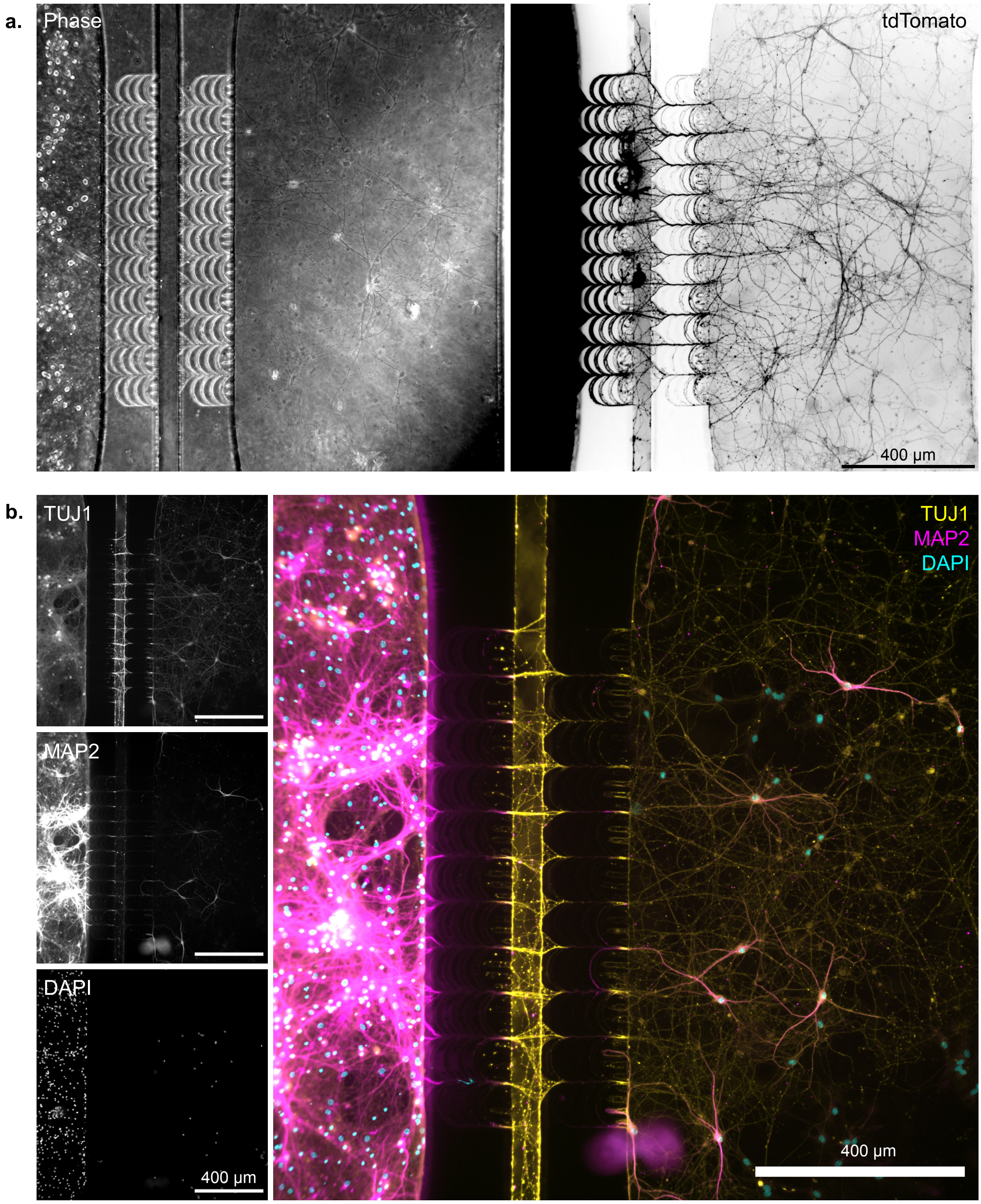


**Supplementary Figure 2**

**Robust axonal invasion from the presyn to the postsyn compartment**

**(a)** Epifluorescence microscopy field of a representative $Hip^{mTmG +/-}$>$Hip^{mTmG -/-}$ network at DIV19. Presynaptic neurons densely innervate the region in front of microchannels in the postsynaptic compartment. **(b)** Epifluorescence microscopy field of a representative Hip>Hip network at DIV24.

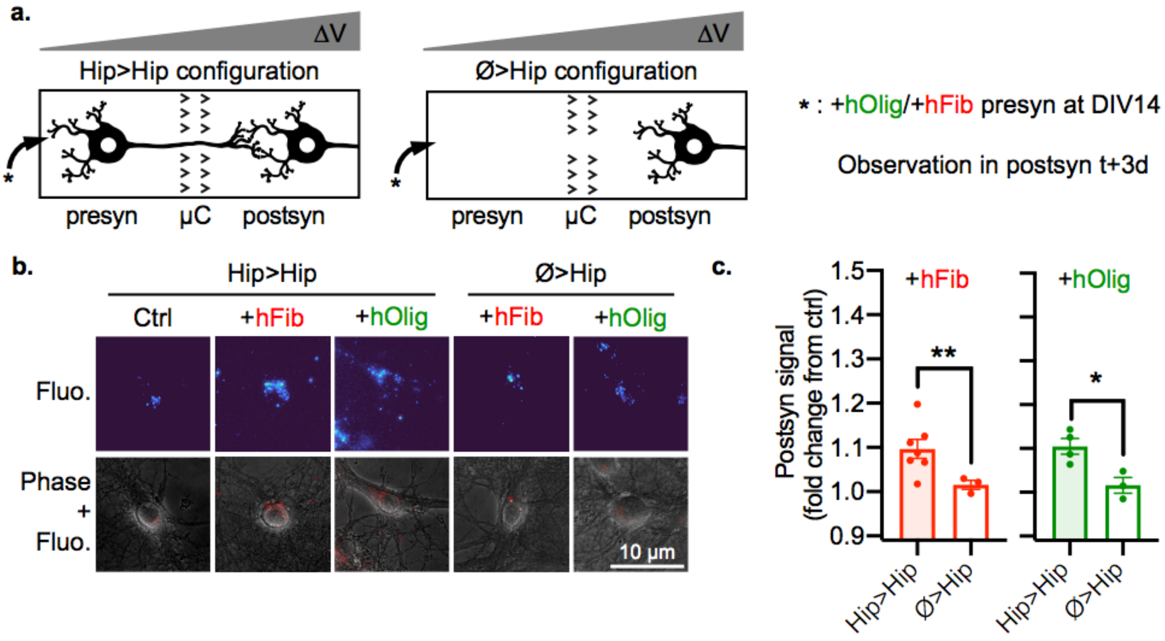


**Supplementary Figure 3**

**aSyn exogenous aggregates are actively transferred from neuron to neuron in an anterograde fashion.**

**(a)** Schematic representation of the experimental design. The presynaptic compartment of Hip>Hip or Ø>Hip networks was spiked with 500 nM hFib or 500 nM hOlig, and aggregates associated fluorescence was monitored in neuronal somas in the postsyn compartment 3 days later. **(b)** Representative epifluorescence microscopy fields of postsynaptic neurons. Top row, hFib associated fluorescence is shown with the Turbo colormap (dark blue is zero, dark red is max. Obtained from github.com/cleterrier/ChrisLUTs.). Bottom row, hFib fluorescence is in red and Phase signal in gray. **(c)** Quantification of the average fluorescence in the somas of postsynaptic neurons (fold change from signal in control condition). hFib networks : n=9-28, N=4-7. hOlig networks : n=10-16, N=3-4. (n individual culture devices from N individual experiments). Unpaired t-test with Welch's correction. Fluo.: aggregates associated fluorescence. Error bars show standard error of the mean.

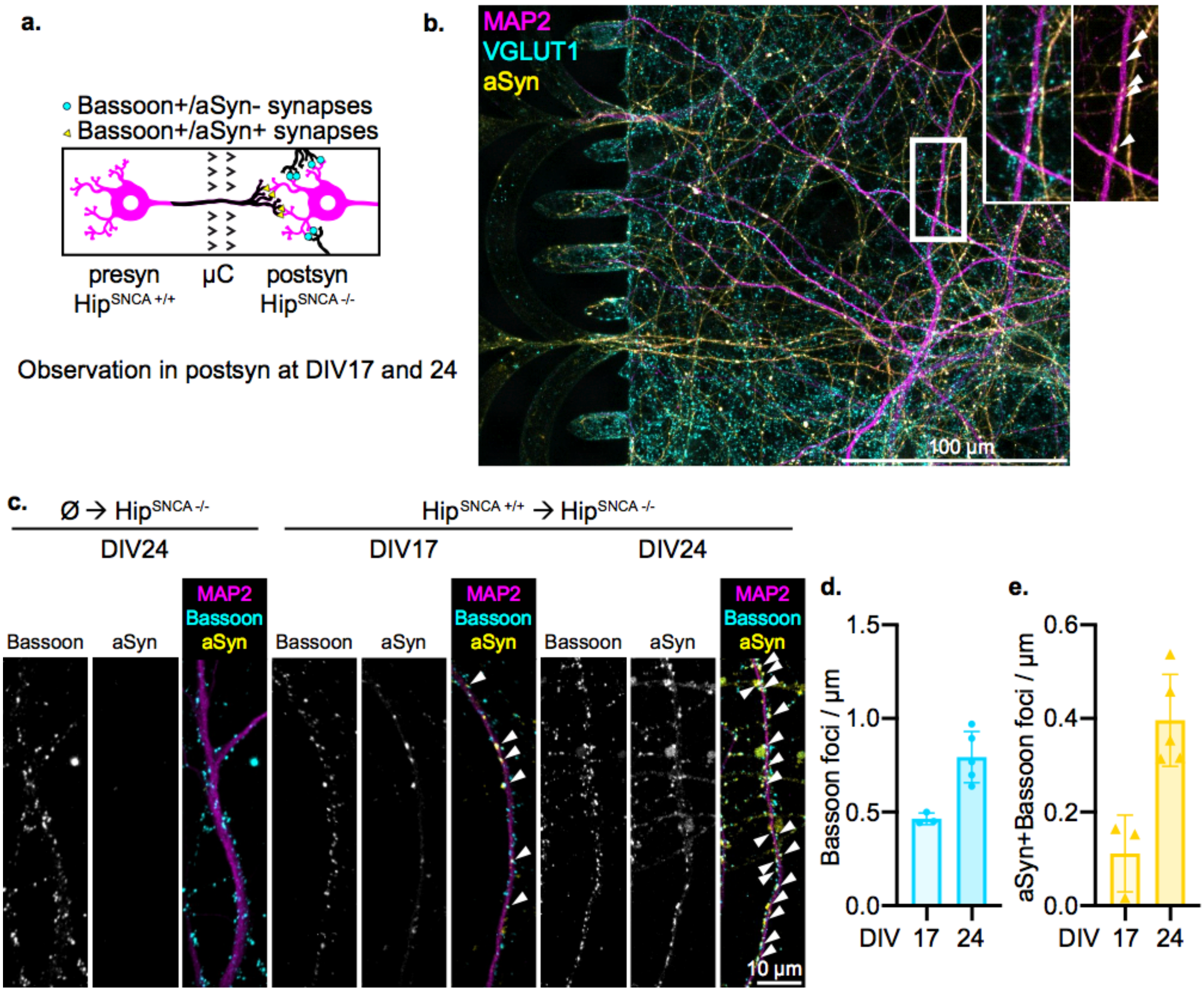


**Supplementary Figure 4**

**Synaptic connectivity between the presynaptic and postsynaptic populations increases with culture time.**

**(a)** Schematic representation of the experimental design. Hip$^{SNCA +/+}$>Hip$^{SNCA -/-}$ networks permitted the estimation of overall synaptic density by staining for synaptic proteins, and of inter-compartment synaptic structures by staining for the presynaptic aSyn protein. **(b)** Representative confocal microscopy field of neurons in the postsynaptic chamber of a 21 days old culture. White arrows highlight synaptic foci stained with both aSyn and VGLUT1. **(c,d,e)** Evolution of synaptic connectivity over culture time. **(c)** Representative confocal microscopy field of dendrites from the postsynaptic compartment of Ø>Hip$^{SNCA -/-}$ and Hip$^{SNCA +/+}$>Hip$^{SNCA -/-}$ networks. **(d)** Quantification of the number of Bassoon foci per µm of dendrites in the postsynaptic compartment. Individual data points represent individual culture devices. **(e)** Quantification of the number of Bassoon foci also positive aSyn per µm of dendrites in the postsynaptic compartment. Individual data points represent individual culture devices. n=3-5 individual culture devices from N=1 individual experiment. Error bars show standard deviation.

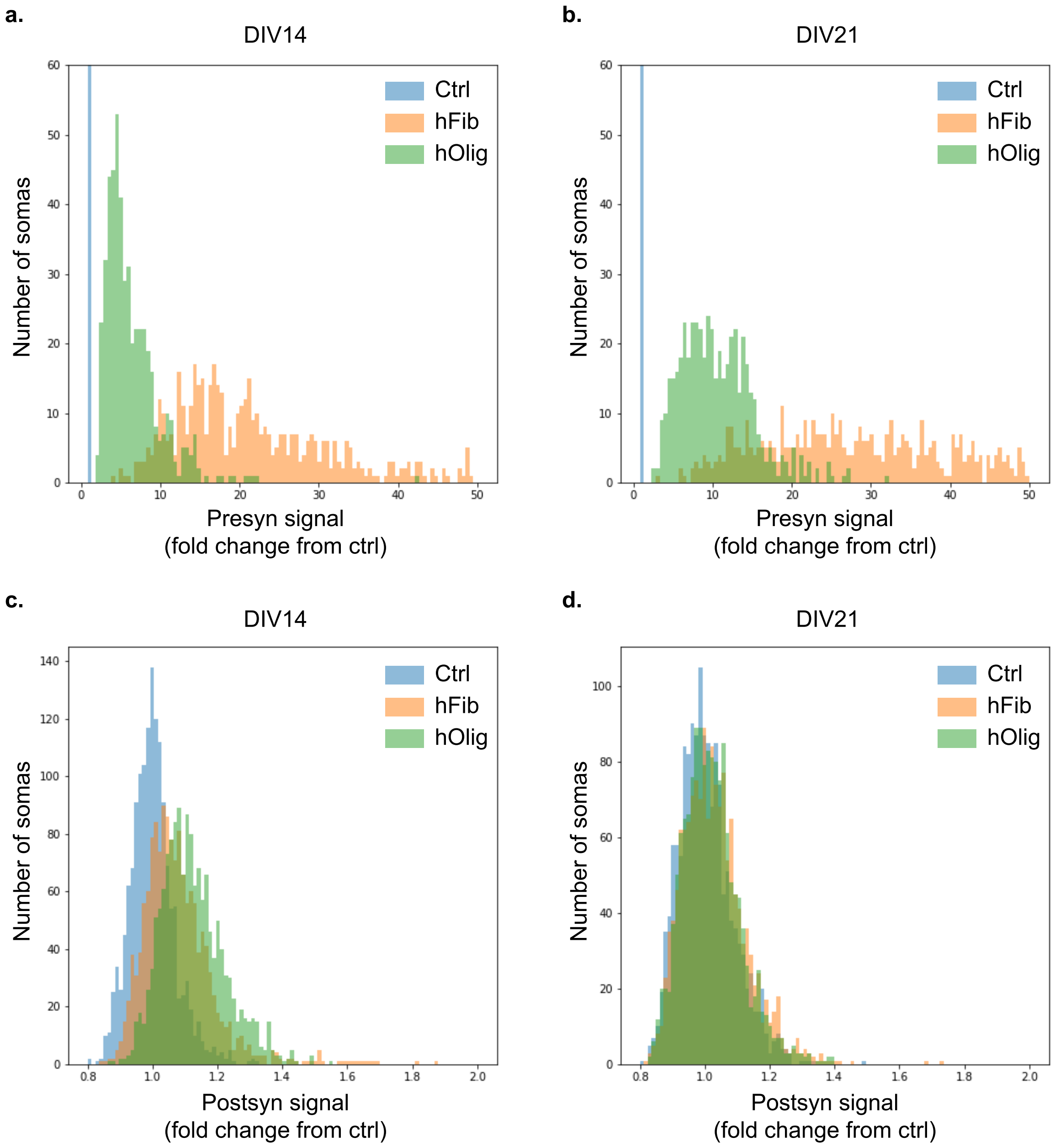


**Supplementary Figure S5**

**Distribution of hFib and hOlig associated fluorescence in presynaptic and postsynaptic neurons.**

Histograms of the distribution of aggregates associated fluorescence in 500 somas randomly picked from n=8-21 individual culture devices from N=2-5 individual experiments. Somas located in the **(a,b)** presynaptic or **(c,d)** postsynaptic compartment of Hip>Hip networks treated at **(a,c)** DIV14 or **(b,d)** DIV21 with control solution (blue), 500 nM of hFib (orange) or 500 nM holig (green).

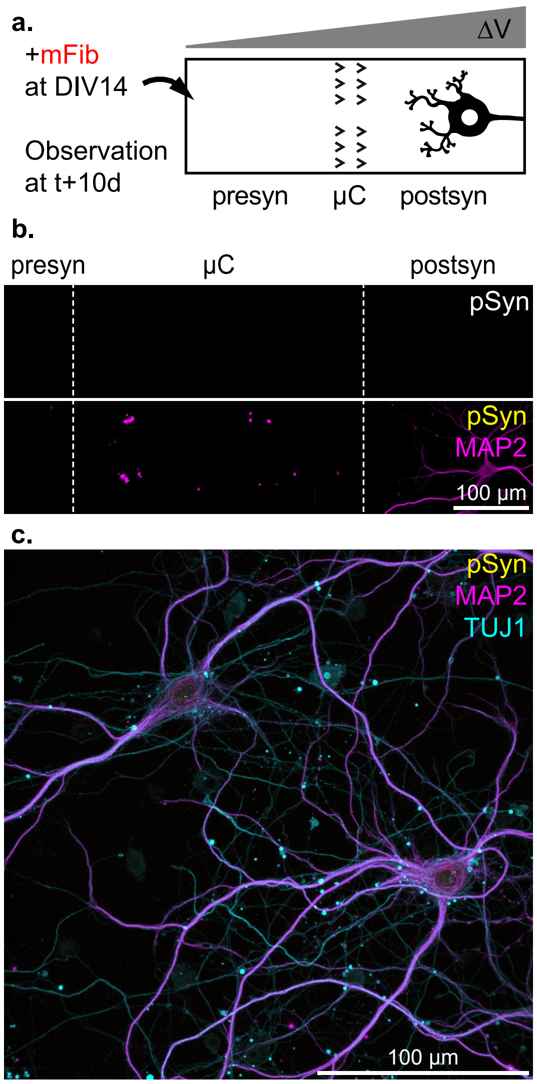


**Supplementary Figure S6**

**Seeding aggregates do not passively diffuse between the culture compartments.**

**(a)** Schematic representation of the experimental design. The presynaptic compartment of Ø>Hip networks was spiked at DIV14 with 500 nM of mFib, and endogenous aSyn aggregation was monitored 10 days later. **(b)** Representative epifluorescence field of a Ø>Hip network. **(c)** Representative confocal microscopy field of the postsynaptic compartment of a Ø>Hip network.
